## Supplementary figures and images for "High-Throughput, Single-Copy Sequencing Reveals SARS-CoV-2 Spike Variants Coincident with Mounting Humoral Immunity during Acute COVID-19"

### Figure S1

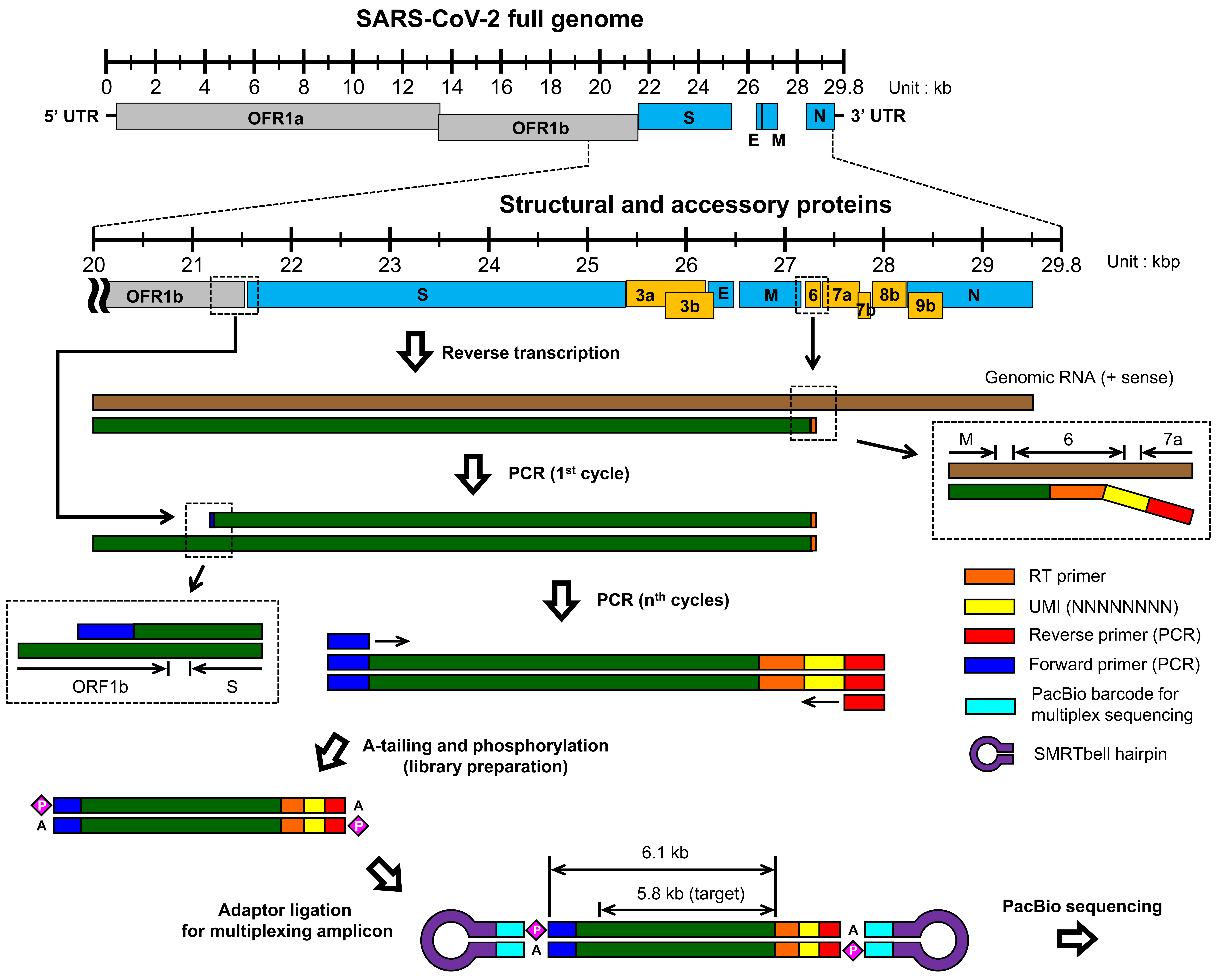

### Figure S2

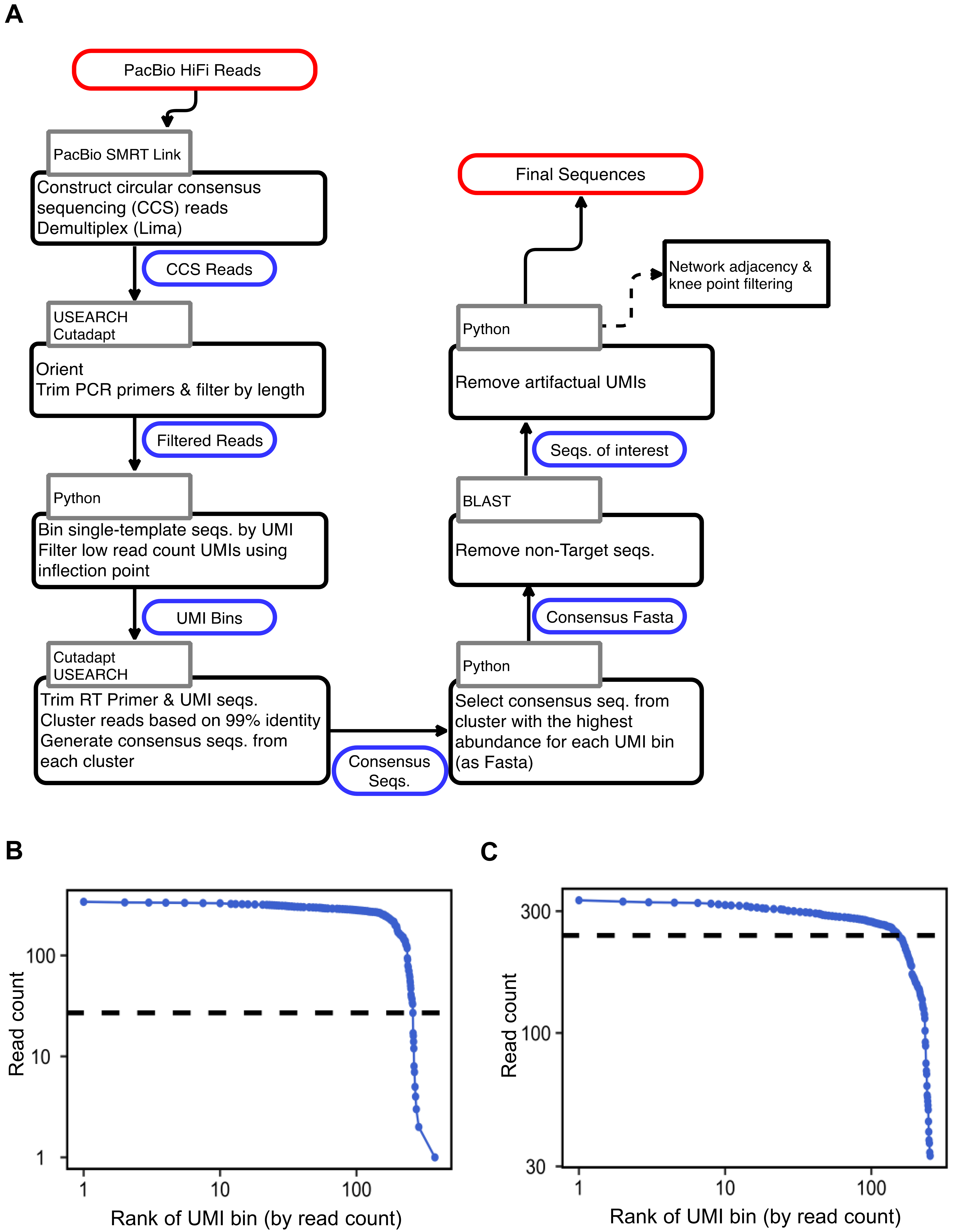

### Figure S3

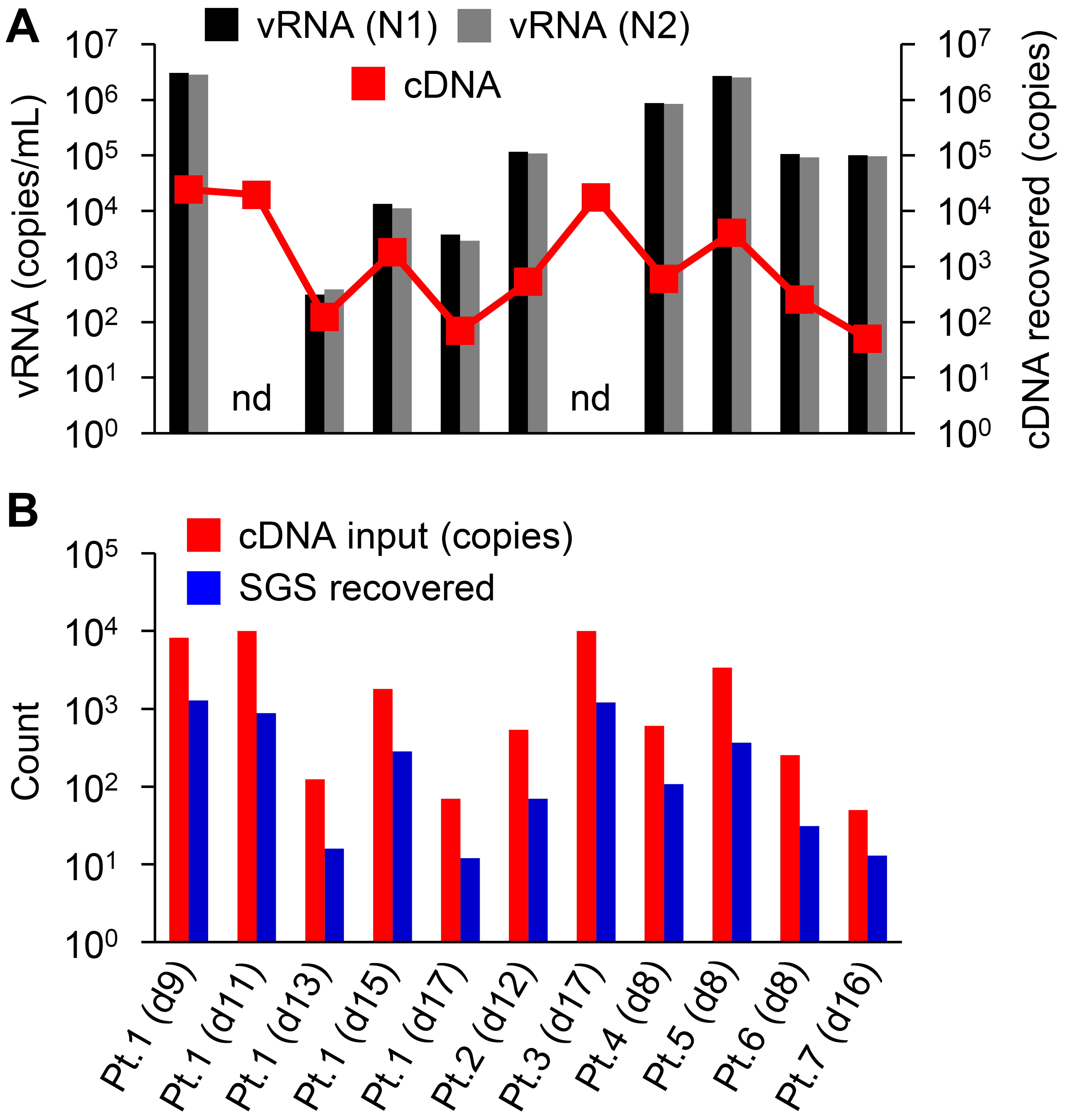

### Figure S4

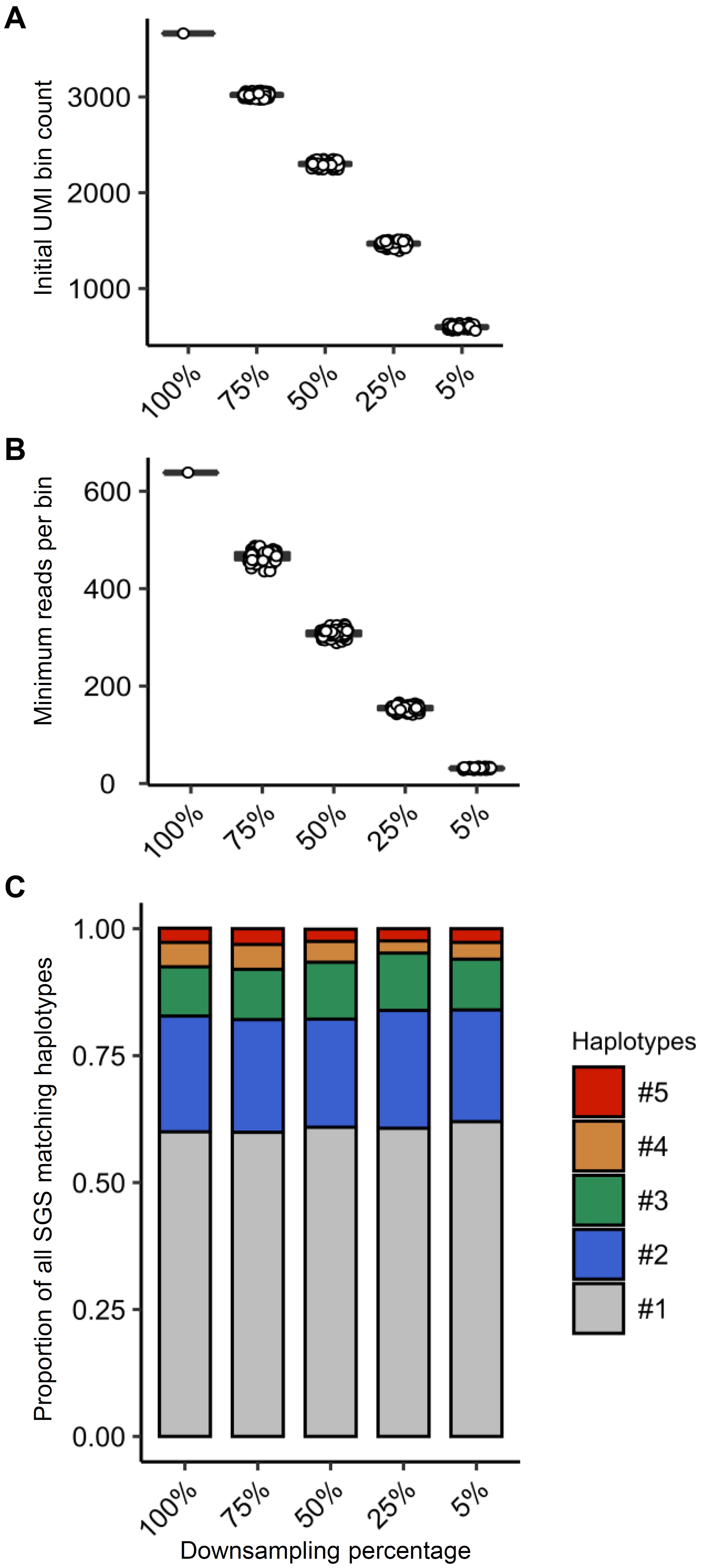
