## Supplementary material for "High-Throughput, Single-Copy Sequencing Reveals SARS-CoV-2 Spike Variants Coincident with Mounting Humoral Immunity during Acute COVID-19": Table S1

| *ID* | *Age* | *Sex* | *Race/ Ethn.* | *Sample Day** | *Comorbidity* | *Suppl. Oxygen* | *Pharmacotherapy* | *Outcome* |
|---|---|---|---|---|---|---|---|---|
| *1* | 38 | F | H | 9, 11, 13, 15, 17 | LCL, HSCT | None | None | Recovered |
| *2* | 60 | M | AA | 12 | Chordoma | None | None | Recovered |
| *3* | 75 | M | H | 17 | Dyslipidemia | MV | Tocilizumab Steroids | Died |
| *4* | 54 | M | W | 8 | Asthma, CLL | MV | Remdesivir | Recovered |
| *5* | 67 | M | AA | 8 | Obesity | MV | Steroids Remdesivir | Died |
| *6* | 22 | M | W | 8 | None | HFNC | None | Recovered |
| *7* | 62 | M | Asian | 16 | GERD, OSA | MV | None | Recovered |

*Sample day denotes day of clinical illness.
H, Hispanic; AA, African-American; W, white; LCL, large cell lymphoma; HSCT, hematopoietic stem cell transplant; CLL, chronic lymphocytic leukemia; GERD, gastro-esophageal reflux disease; OSA, obstructive sleep apnea; MV, mechanical ventilation; HFNC, high-flow nasal cannula; LOS, length of stay.
