## Supplementary material for "High-Throughput, Single-Copy Sequencing Reveals SARS-CoV-2 Spike Variants Coincident with Mounting Humoral Immunity during Acute COVID-19": Table S2

| Primer set | Primer name | MN908947 coordinates (GenBank) | Sequence (5’ → 3’) |
|---|---|---|---|
| Reverse transcription | SEM-RT | 27357-27379 | CCGCTCCGTCCGACGACTCACTATANNNNNNNNTCTCCATTGGTTGCTCTTCATCT |
| FCA forward | FCA_F | 22195-22216 | TTTAGTGCGTGATCTCCCTCAG |
| FCA reverse | FCA_R | 22588-22608 | GATGCAAATCTGGTGGCGTTA |
| PCR forward-1 | PCR_F1 | 21217-21240 | TTCGCATGGTGGACAGCCTTTGTT |
| PCR forward-2 | PCR_F2 | 21288-21311 | TTATCTTGGCAAACCACGCGAACA |
| PCR reverse | PCR_R | Universal primer | CCGCTCCGTCCGACGACTCACTATA |
